# High-fat diets promote peritoneal inflammation and augment endometriosis-associated abdominal hyperalgesia

**DOI:** 10.1101/2023.11.09.566474

**Authors:** Tristin Herup-Wheeler, Mingxin Shi, Madeleine E. Harvey, Chandni Talwar, Ramakrishna Kommagani, James A. MacLean, Kanako Hayashi

## Abstract

Immune dysfunction is one of the central components in the development and progression of endometriosis by establishing a chronic inflammatory environment. Western-style high-fat diets (HFD) have been linked to greater systemic inflammation to cause metabolic and chronic inflammatory diseases, and are also considered an environmental risk factor for gynecologic diseases. Here, we aimed to examine how HFD alter an inflammatory environment in endometriosis and discern their contribution to endometriotic-associated hyperalgesia. Our results showed that HFD-induced obesity enhanced abdominal mechanical allodynia that was induced by endometriotic lesions. Peritoneal inflammatory macrophages and cytokine levels increased by lesion induction were elevated by chronic exposure to HFD. Pain-related mediators in the dorsal root ganglia were further stimulated after lesion induction under the HFD condition. Although HFD did not affect inflammatory macrophages in the peritoneal cavity without lesion induction, the diversity and composition of the gut microbiota were clearly altered by HFD as a sign of low-grade systemic inflammation. Thus, HFD alone might not establish a local inflammatory environment in the pelvic cavity, but it can contribute to further enhancing chronic inflammation, leading to the exacerbation of endometriosis-associated abdominal hyperalgesia following the establishment and progression of the disease.

## Introduction

Endometriosis is a chronic and incurable inflammatory disorder and affects approximately 10% of reproductive-aged women (1, 2). It is associated with debilitating chronic pelvic pain and infertility, which substantially reduce the quality of life of women and their families (3, 4). Because endometriosis is estrogen-dependent, current treatments focus on inhibiting estrogen production and function. However, hormonal treatments and surgical excision of lesions are often of limited efficacy with high recurrence rates, frequent side effects, additional costs, and potential morbidity (5). As nearly 70% of patients suffer unsolved chronic pain and other related conditions (6), the direct costs of endometriosis were estimated at $12,118 per patient per year in the US, and indirect costs were $15,737 (7). The pathogenesis of endometriosis is a complex process and remains to be fully understood. Retrograde menstruation has been widely accepted as the origin of endometriotic tissues (8). However, as retrograde menstruation occurs in more than 90% of menstruating women (9), the pathogenesis of the disease is not well understood, and other factors must contribute to establishing endometriotic lesions and disease progression (1, 4, 10).

Obesity is an epidemic health burden affecting nearly 40% of adults and 18% of children in the United States (11). Being overweight and obese are considered critical risk factors for chronic diseases, as fat accumulation causes low-grade chronic inflammation (12) characterized by immune cell infiltration into adipose tissues and elevated proinflammatory factors (13). Moreover, excessive fat consumption and accumulation in the body alter gut microbiota, resulting in dysbiosis to induce low-grade systemic inflammation (14). Obesity-induced inflammation is associated with metabolic and autoimmune disorders in women, causing reproductive dysfunctions such as polycystic ovary syndrome (PCOS), implantation and pregnancy failure, and pregnancy complications, including miscarriages (15–18). While endometriosis is a chronic inflammatory disease, several epidemiological studies have reported an inverse correlation between endometriosis and body mass index (BMI) (19). However, obesity does not protect against endometriosis (19), and BMI is correlated with the severity but not the frequency of disease diagnoses (20). Thus, BMI does not provide a simple risk factor for a heterogeneous endometriotic disease as it does not consider different components of excess weight, such as adipose deposit location and interaction with neighboring tissues (20, 21). Additionally, the correlation between diet-induced obesity and endometriosis-associated pain or hypersensitivity, one of the significant endometriosis symptoms, has not been addressed.

Rigorous prior research suggests that aberrant inflammation contributes to the onset and progression of endometriosis (22–27). Macrophages (MΦ) are considered to be key players in promoting disease progression (25, 28, 29), as abundant MΦ are present in ectopic lesions (30) and elevated in the pelvic cavity (31). These MΦ populations establish an inflammatory environment in the pelvic cavity by secreting cytokines and chemokines, which encourage lesion growth and progression (24, 28, 29, 32, 33) and contribute to endometriosis-associated pelvic pain (32, 34, 35). Diet-induced obesity dysregulates immune cells to induce cytokine secretion (13, 36, 37), increasing the risks of chronic pain. Therefore, the present study seeks to understand whether high-fat diets (HFD) affect the progression of endometriosis disease and immune dysfunctions and how HFD influences endometriosis-associated hyperalgesia.

The present results highlight that endometriosis-associated abdominal hyperalgesia was escalated in lesion-induced HFD mice according to the results of the behavior study and elevated pain-related mediators in the dorsal root ganglion (DRG). Increased proinflammatory MΦ (Ly6C+ MΦ) and cytokines by lesion induction were further enhanced by exposure to HFD. The results also indicate that gut microbiota dysbiosis under the HFD condition contributed to an aberrant inflammatory environment and sensitized endometriosis-associated hypersensitivity.

## Results

### Diet-induced Obesity on endometriosis in mice

To examine the effect of diet-induced obesity on endometriosis, female mice were fed HFD containing 45% fat by calories or standard diets (SD) from the age of 5 weeks (defined as Week 0 of the 18-week program, Figure 1A). We chose to start the study at the age of 5 weeks, as this is the adolescent age of mice, corresponding to the teenage period for humans. Mice on the 45% fat diets become obese and are considered physiologically similar to the typical Western diets that contain 36-40% fat by energy (38, 39). A standard rodent diet contains approximately 10% fat (38, 40), similar to a regular diet at Washington State University, which contains 9% of total calories from fat. As shown in Figure 1BCD, body weight (BW), blood glucose, and plasma insulin levels were significantly increased in the group of HFD with/without lesion induction. As baselines, we assessed glucose and insulin levels at 12 weeks after SD or HFD feeding. As previously reported (41, 42), we observed similar BW, glucose, and insulin increases in the HFD group. However, lesion numbers were not altered by HFD compared with SD at 6 weeks after endometriosis-like lesions (ELL) induction (Figure. 2A). We have previously reported that peritoneal MΦ or monocytes are infiltrated into the ELL (28). We thus addressed MΦ infiltration in the lesion staining with CD68, a macrophage marker. CD68+ MΦ were significantly increased within lesions from mice in the HFD group (Figure. 2BC), indicating MΦ infiltration was accelerated in the ELL-HFD mice, although this did not appear to affect lesion development.

**Figure. 1.**
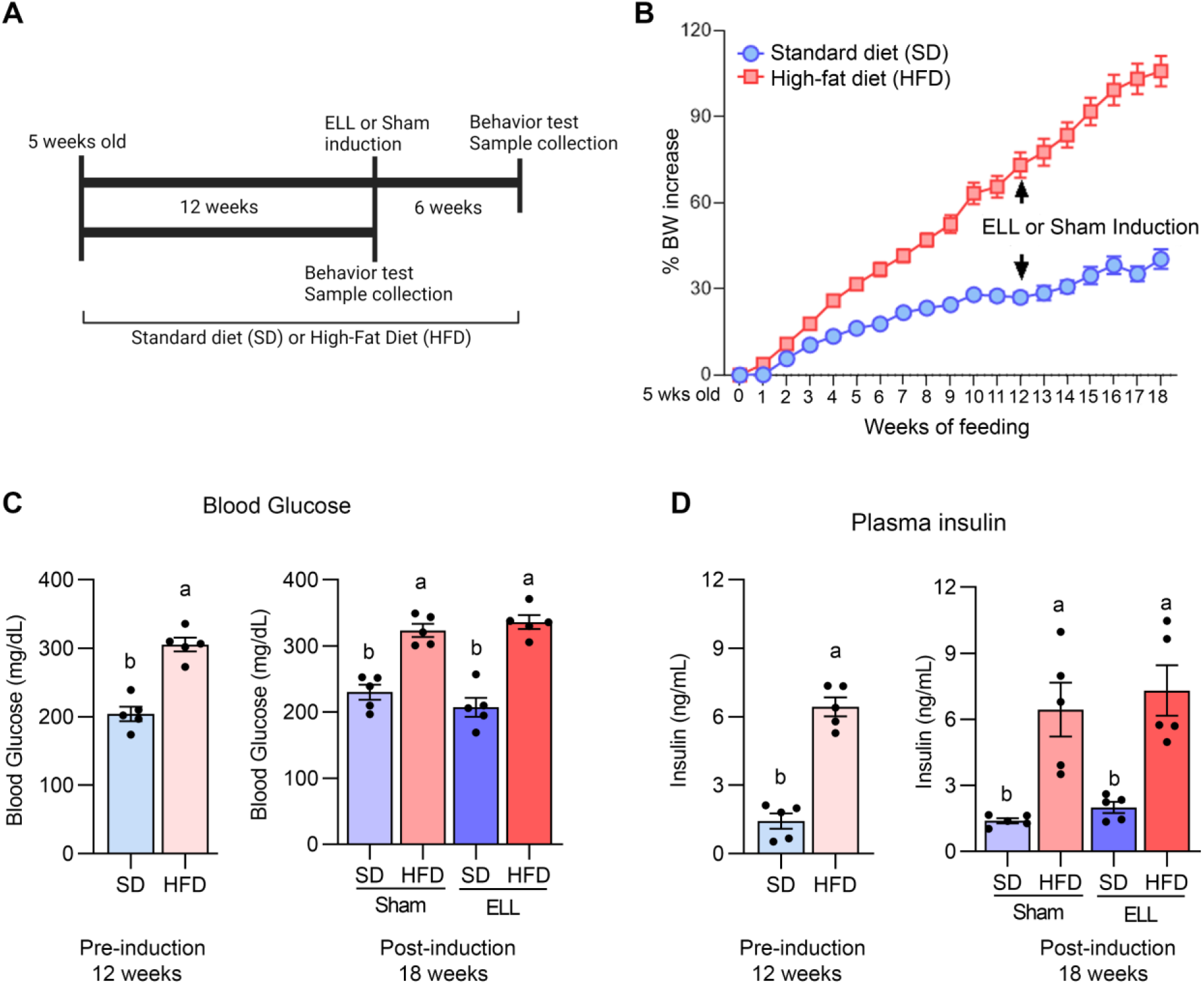
Diet-induced obesity in the mouse model of endometriosis. (**A**) Experimental study design as described in Material and Methods. (**B**) Body weight (BW) changes in mice during the feeding of standard diets (SD) or 45% high-fat diets (HFD). Female mice were fed either SD or HFD starting at the age of 5 weeks (defined as Week 0). (**C**) Blood glucose levels by cardiac puncture were measured by Contour Next (n=5). (**D**) Plasma insulin levels were quantified by ELISA (n=5). Data at 12 weeks were analyzed by two-tailed Student’s t-test comparing SD and HFD. Data at 18 weeks were analyzed through one-way ANOVA and Tukey’s post hoc test. Values in graphs are expressed as the mean ± SEM. Statistical differences among the groups were reported with the compact letter display (shown as a vs b; P<0.05). ELL: endometrial-like lesion.

**Figure. 2.**
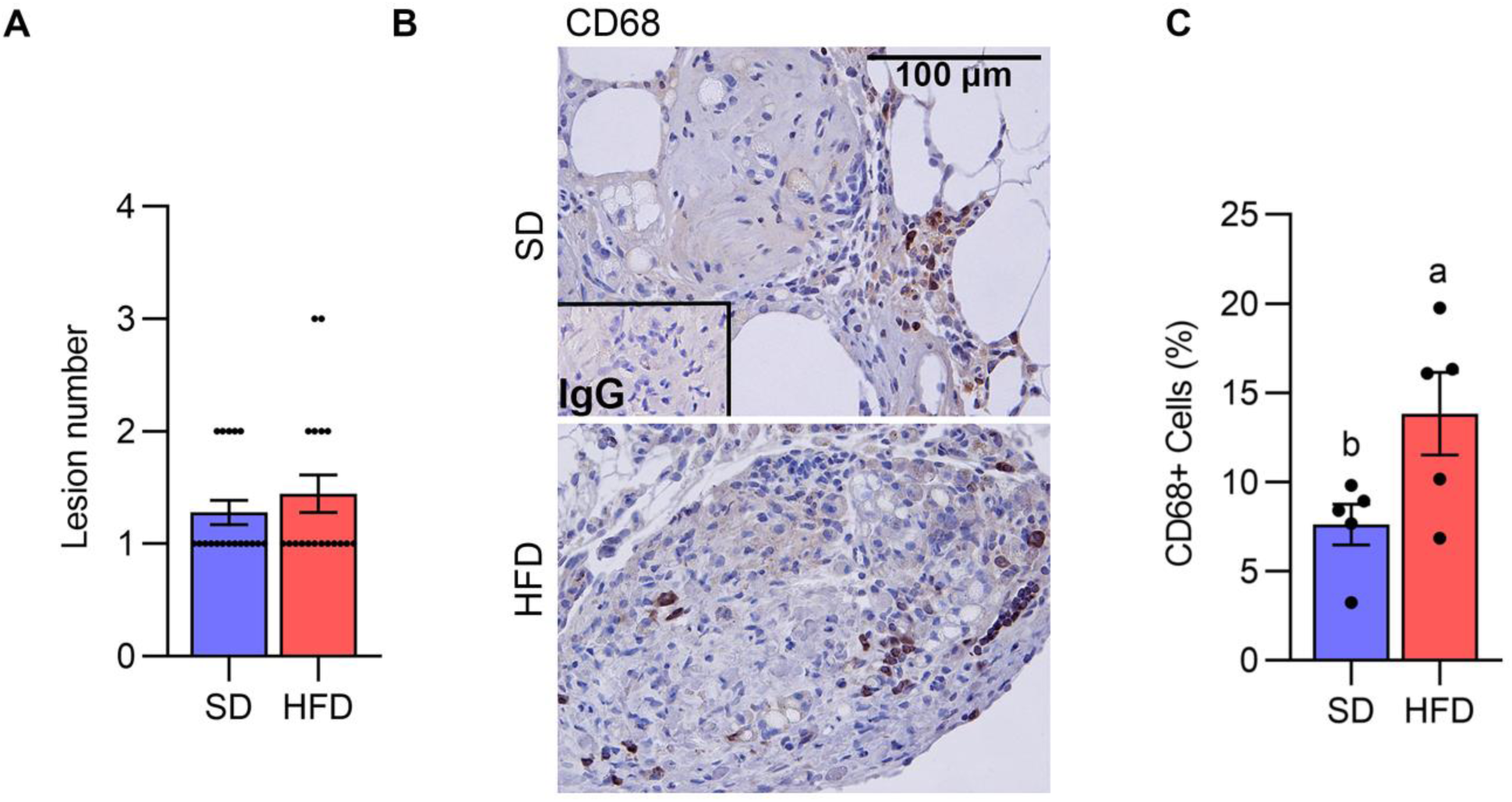
Diet-induced obesity increases macrophage infiltration in the lesion. (**A**) Lesion number (n=18). (**B**) CD68 was stained to determine macrophage infiltration in the lesion. (**C**) The quantification of the percentage of CD68+ cells per total cells (n=5). Data were analyzed with the student t-test and are shown as mean ± SEM. a vs b; P<0.05. SD: standard diets, and HFD: high-fat diets.

### HFD accelerates endometriosis-associated abdominal hyperalgesia

Since HFD can induce chronic pain, we next performed the von Frey test to examine the abdominal and hind paw retraction threshold to determine whether HFD affects endometriosis-associated hyperalgesia. To establish the baseline behavior of our mice, we assessed the abdominal and hind paw retraction threshold at 12 weeks after SD or HFD feeding. The pre-induction stage did not alter the abdominal and hind paw retraction threshold (Figure. 3AC).

**Figure. 3.**
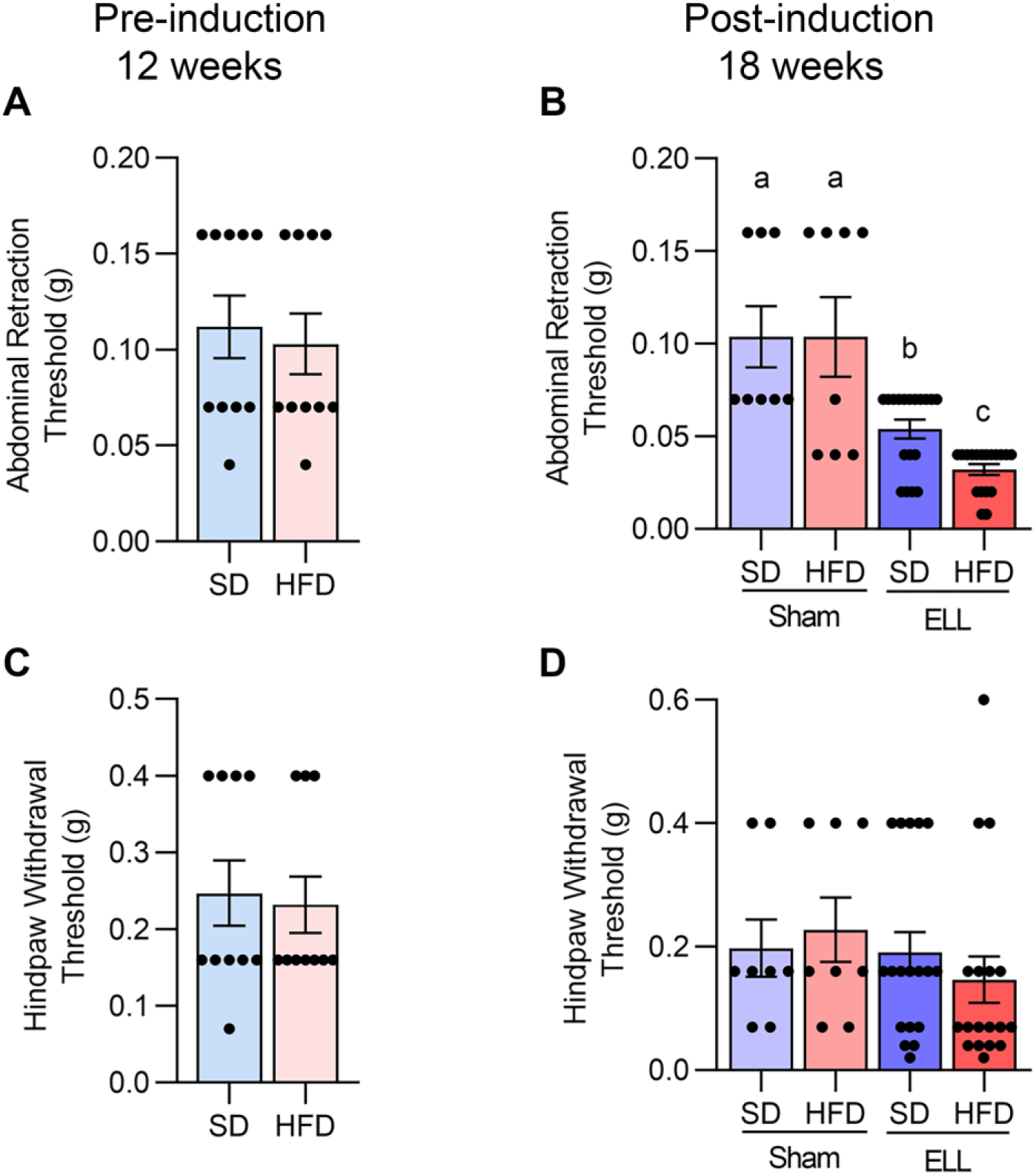
HFD accelerates endometriosis-associated abdominal hyperalgesia. Von Frey tests were performed on mice to the lower abdomen and hind paw at the pre-induction time point after 12 weeks of SD or HFD feeding (**A** and **C**, n=10), or 6 weeks post-lesion induction (**B** and **D**, a total of 18 weeks of SD or HFD feeding, n=8 for Sham and n=18 for ELL groups). Data are shown as mean ± SEM. Statistical significance was determined by student t-test (**A** and **C**), or one-way ANOVA followed by Tukey’s post hoc test (**B** and **D**). a vs b; P<0.05. ELL: endometrial-like lesion, SD: standard diets, and HFD: high-fat diets.

Before examining the effect of HFD on endometriosis-associated hyperalgesia, we first examined how lesion induction time-dependently alters endometriosis-associated hyperalgesia in the mouse model. Three days after ELL induction, mice withdrew both abdominal and hind paw retraction thresholds with significantly lighter stimuli compared with those before ELL induction on Day -1 (Supplementary Figure. S1AB). The abdomen and hind paw retraction sensitivity continued until 3 weeks after ELL induction. By Day 42, 6 weeks after ELL induction, the hind paw retraction threshold was no longer significantly different from Day -1, indicating that systemic peripheral hyperalgesia gradually recovered, whereas the local abdomen was still sensitive. Since we examined the effect of chronic HFD exposure on endometriosis-associated hyperalgesia, we chose a chronic stage, 6 weeks after ELL induction, for further analysis, as endometriosis is a chronic disease, and most patients suffer chronic pelvic pain. Furthermore, the timing of disease onset in endometriosis is currently impossible to determine in patients, and the disease diagnosis typically relies on the woman noticing chronic symptoms.

The abdominal and hind paw sensitivity with SD or HFD were evaluated 6 weeks after lesion induction (18 weeks of SD or HFD feeding). As expected, a significant difference was observed in the abdominal retraction threshold between Sham and ELL, with ELL mice withdrawing from lighter stimuli than Sham mice (Figure. 3B). Importantly, ELL-HFD mice were more sensitive than ELL-SD mice (Figure. 3B). On the other hand, we did not observe any differences in hind paw retraction threshold among the post-induction groups (Figure. 3D).

### HFD increases Ly6C+ MΦ in the peritoneal fluid (PF) of ELL mice

As we observed increased MΦ infiltration in the lesions of the HFD group (Figure. 2BC) and ELL-HFD mice have increased hypersensitivity in the abdomen (Figure. 3B), we expected to observe differences in the inflammatory environment that is established in the peritoneal cavity. Therefore, we assessed immune cell profiles, MΦ, B- and T-cells, in the peritoneal cavity (Figure. 4 and Supplementary Figure. S2). CD11b+ MΦ, CD3+ T-cells, and CD19+ B-cells were not altered by either HFD feeding or lesion induction (Figure. 4AB). We have previously reported that the presence of ELL enhanced the differentiation of recruited (=proinflammatory Ly6C+) MΦ and increased the ablation of embryo-derived resident MΦ (TIM4+ MΦ) (29). We thus examined Ly6C+ cells (monocytes and MΦ), Ly6C+ MΦ, and TIM4+ MΦ. High levels of Ly6C+ cells and Ly6C+ MΦ were observed in the ELL-HFD mice (Figure. 4ABC). In particular, Ly6C+ MΦ were further increased in the ELL-HFD mice than those in ELL-SD mice (Figure. 4C). In agreement with our previous study (29), TIM4+ MΦ were reduced in ELL-SD and ELL-HFD mice (Figure. 4C). Ly6C+ cells, Ly6C+ MΦ and TIM4+ MΦ, as well as CD11b+ MΦ, CD19+ B-cells, and CD3+ T-cells were not affected by HFD feeding at the pre-induction stage (Supplementary Figure. S2). These results suggest that ELL induction under the HFD condition further increases proinflammatory Ly6C+ MΦ in the peritoneal cavity.

**Figure. 4.**
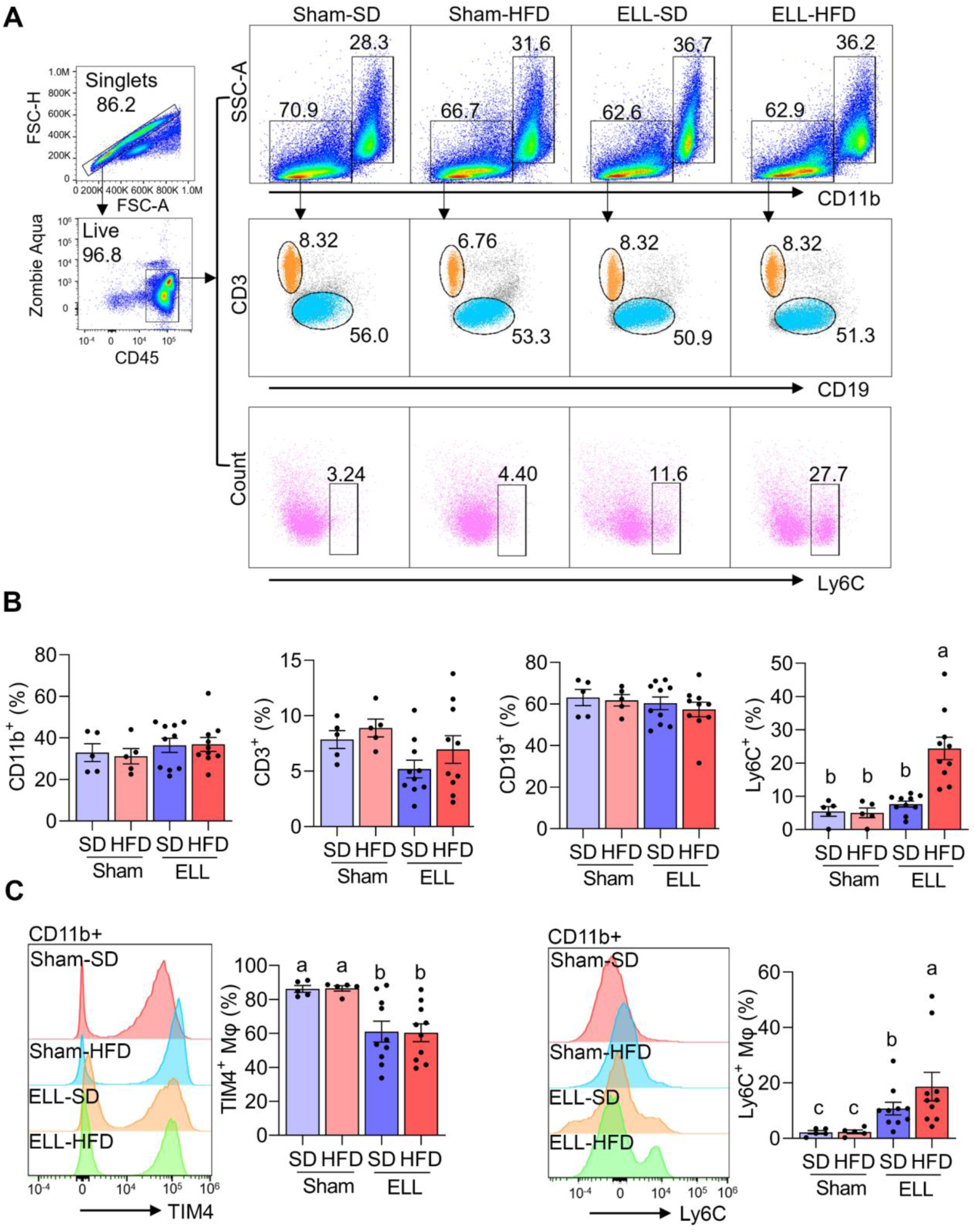
HFD increases Ly6C+ macrophages (MΦ) in the peritoneal fluid (PF) of ELL mice. (**A**) Flow cytometer analysis for CD11b+ (MΦ), CD3+ (T-cells), CD19+ (B-cells), and Ly6C+ (monocytes and MΦ) cells in the PF. (**B**) Quantification of CD11b+, CD3+, CD19+, and Ly6C+ cells in the groups of Sham-SD (n=5), Sham-HFD (n=5), ELL-SD (n=10) and ELL-HFD (n=10). (**C**) TIM4+ and Ly6C+ MΦ were quantified in the PF. Data were analyzed through One-way ANOVA followed by Tukey’s post hoc test and expressed as the mean ± SEM. a vs b vs c; p<0.05. ELL: endometrial-like lesion, SD: standard diets, and HFD: high-fat diets.

### HFD altered peritoneal cytokines in the ELL mice

Proinflammatory MΦ secrete cytokines, chemokines, and growth factors that establish the inflammatory environment (27, 43). Abundant cytokines and chemokines have been observed in the pelvic cavity of endometriosis patients (24, 27). Specifically, the levels of TNFα, IL1β, and IL6 are increased in pelvic MΦ isolated from endometriosis patients (44). Thus, we next examined the secretion of proinflammatory factors, TNFα, IL1β, and IL6, as well as IL10, which is known to possess immunoregulatory function and antiinflammatory properties (Figure. 5A-D). In support of previous reports, TNFα and IL1β levels were elevated in the ELL groups compared with those in the Sham group, while TNFα was further increased in ELL-HFD mice. IL6 tended to be increased by lesion induction in both SD and HFD groups, though we did not see significant differences. IL10 levels were not significantly altered among the groups of Sham-SD, Sham-HFD, and ELL-SD mice, whereas it was significantly lower in the ELL-HFD mice compared with that of ELL-SD mice.

**Figure. 5.**
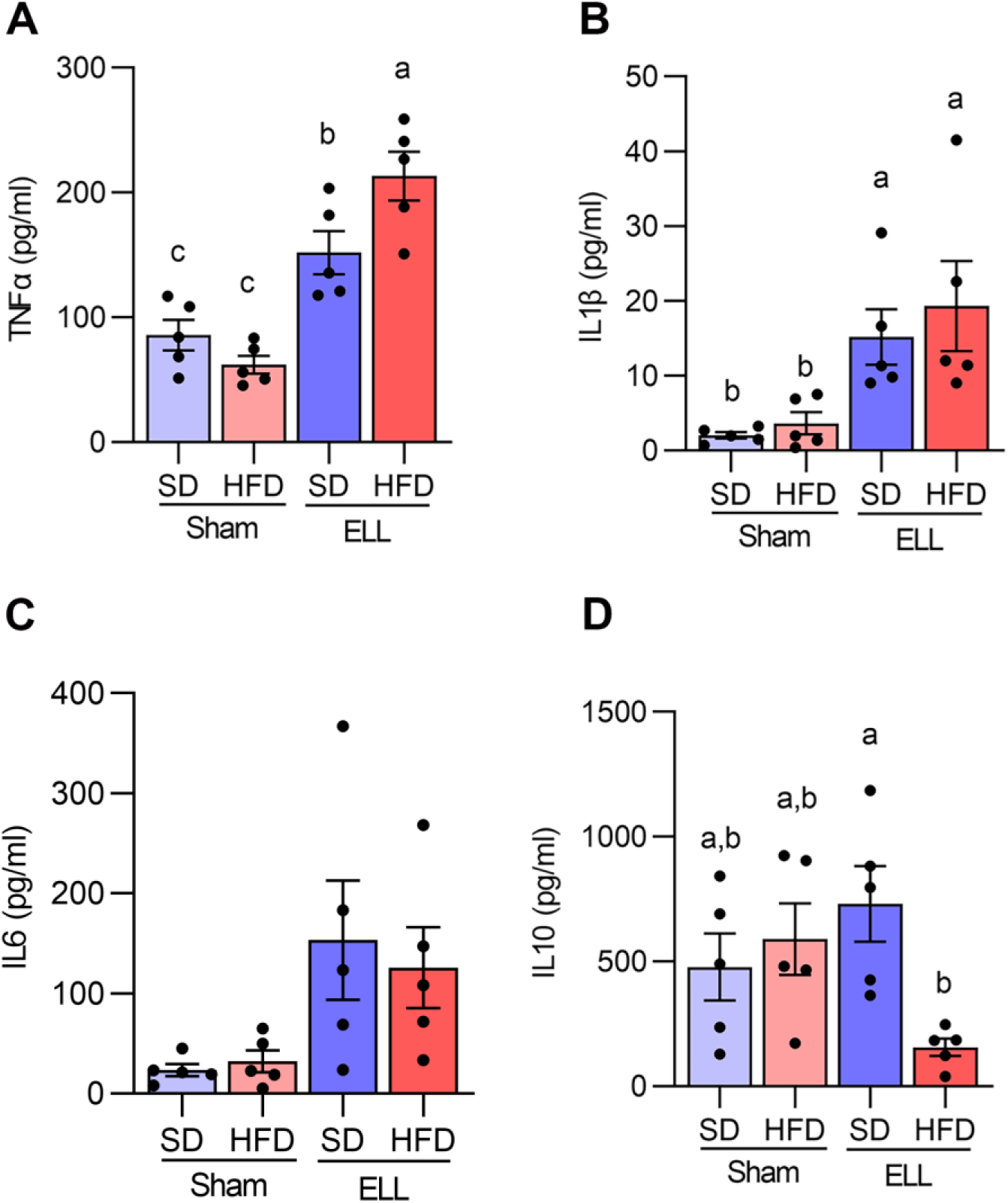
Quantification of TNFα, IL1β, IL6, and IL10 in the peritoneal fluid (PF). Peritoneal (**A**) TNFα, (**B**) IL1β, (**C**) IL6, and (**D**) IL10 were measured with IQELISA and analyzed with ANOVA followed by Tukey’s post hoc test. Values in graphs are expressed as the mean ± SEM (n=5). a vs b vs c; P<0.05. ELL: endometrial-like lesion, SD: standard diets, and HFD: high-fat diets.

### HFD stimulates pain-related mediators in the DRG of ELL mice

Aberrant accumulation of inflammatory factors can stimulate peripheral nerve terminals of nociceptor neurons innervating different tissues in peripheral organs (45), resulting in an increase in the expression of transient receptor potential channels e.g., TRPA1 and TRPV1. Activation of peripheral nerves is also associated with the increased release of neurotransmitters and neuromodulators such as SP, CGRP, and BDNF. BDNF is known to regulate both initiation and maintenance of chronic endometriosis-associated pain (46, 47) involving neuroangiogenesis (48) and innervation in the pelvic organs (45). We thus examined the inflammatory mediators, neurotransmitters, and neuromodulators in the L4-6 DRG, which are the primary spinal ganglia receiving sensory input from pelvic organs (Figure. 6AB). Significantly more BDNF+ neurons were observed in mice fed HFD. BDNF+ neurons were higher in mice when ELL were present and most abundant in the HFD-ELL group. In contrast, CGRP+ neurons were only significantly elevated in the ELL-HFD mice. SP+ neurons were elevated by lesion induction, while HFD further increased SP+ neurons after ELL induction. Although the numbers of TRPV1+ neurons were relatively consistent between Sham- and ELL-mice, there was a significant difference between ELL-HFD mice and ELL-SD mice. These results suggest that lesion induction and/or HFD feeding stimulate endometriosis-associated peripheral pain mediators.

**Figure. 6.**
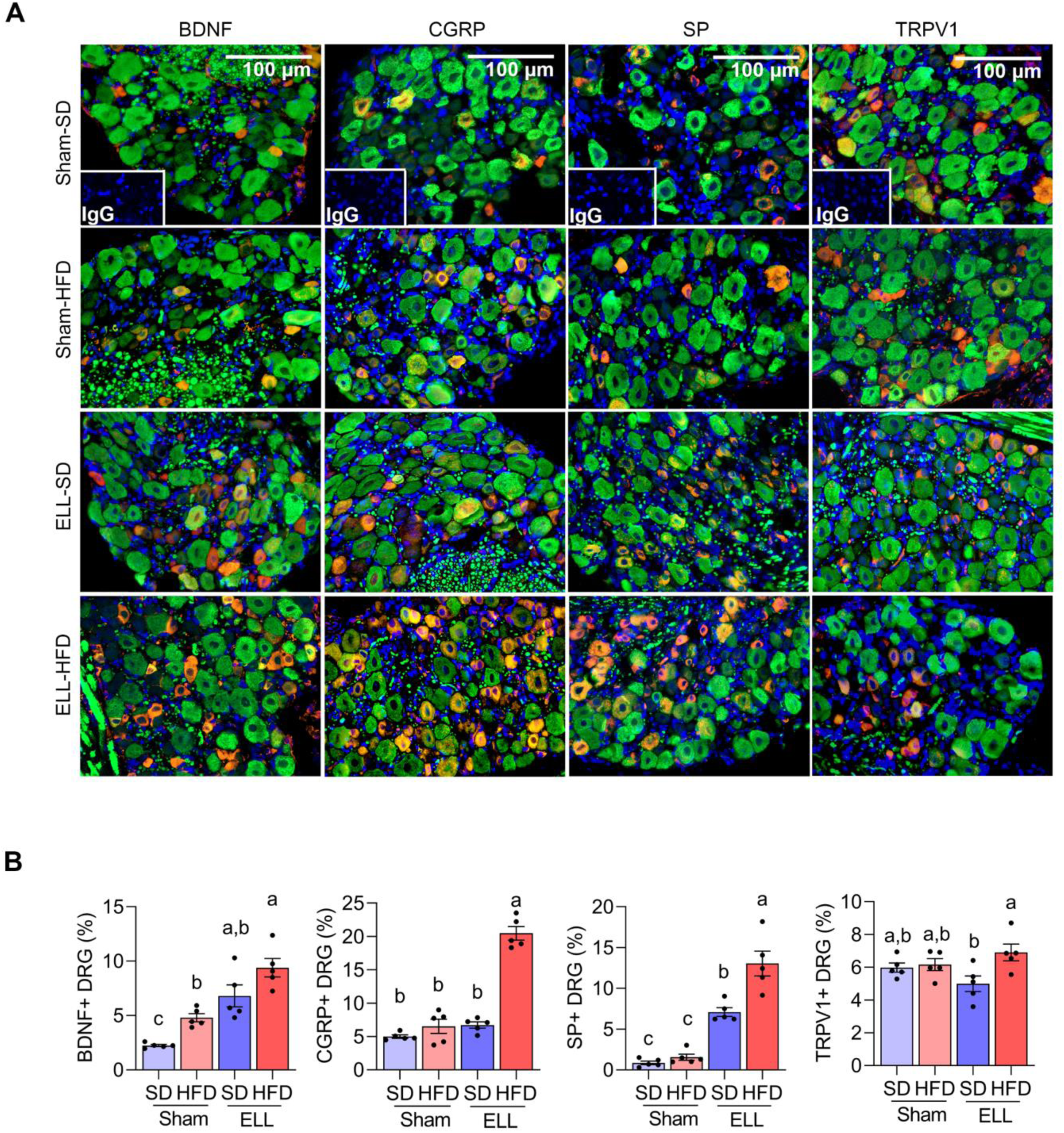
HFD stimulates pain-related mediators in the DRG of ELL mice. (**A**) Immunofluorescence results of BDNF, CGRP, SP, TRPV1, and neurofilament (NF, green) in DRG. NF was used as a marker of DRG cell body and was co-stained with BDNF, CGRP, SP, or TRPV1. (**B**) BDNF, CGRP, SP, or TRPV1 positive DRG per NF positive DRG was counted and quantified (n=5 per group). One-way ANOVA followed by Tukey’s post hoc test was used for statistical analysis. Data were shown as mean ± SEM. a vs b vs c; P<0.05. ELL: endometrial-like lesion, SD: standard diets, and HFD: high-fat diets. DRG: dorsal root ganglia.

### HFD altered the composition of the gut microbiota

As increased fat accumulation alters gut microbiota and causes low-grade systemic inflammation (14), we next examined 16S rRNA gene sequencing of DNA isolated from fecal samples in SD or HFD with/without ELL-induced mice (Figure 7). Microbial alpha diversity was lower in the feces of HFD-fed mice than in SD-fed mice (Figure 7A). Principal coordinates analysis (PCoA) showed uniquely clustered microbial variance induced by HFD (Figure 7B). However, ELL induction did not alter microbial diversity or variance, indicating that long-term systemic alterations induced by HFD affect the composition of the gut microbiota more so than lesion induction in mice.

**Figure. 7.**
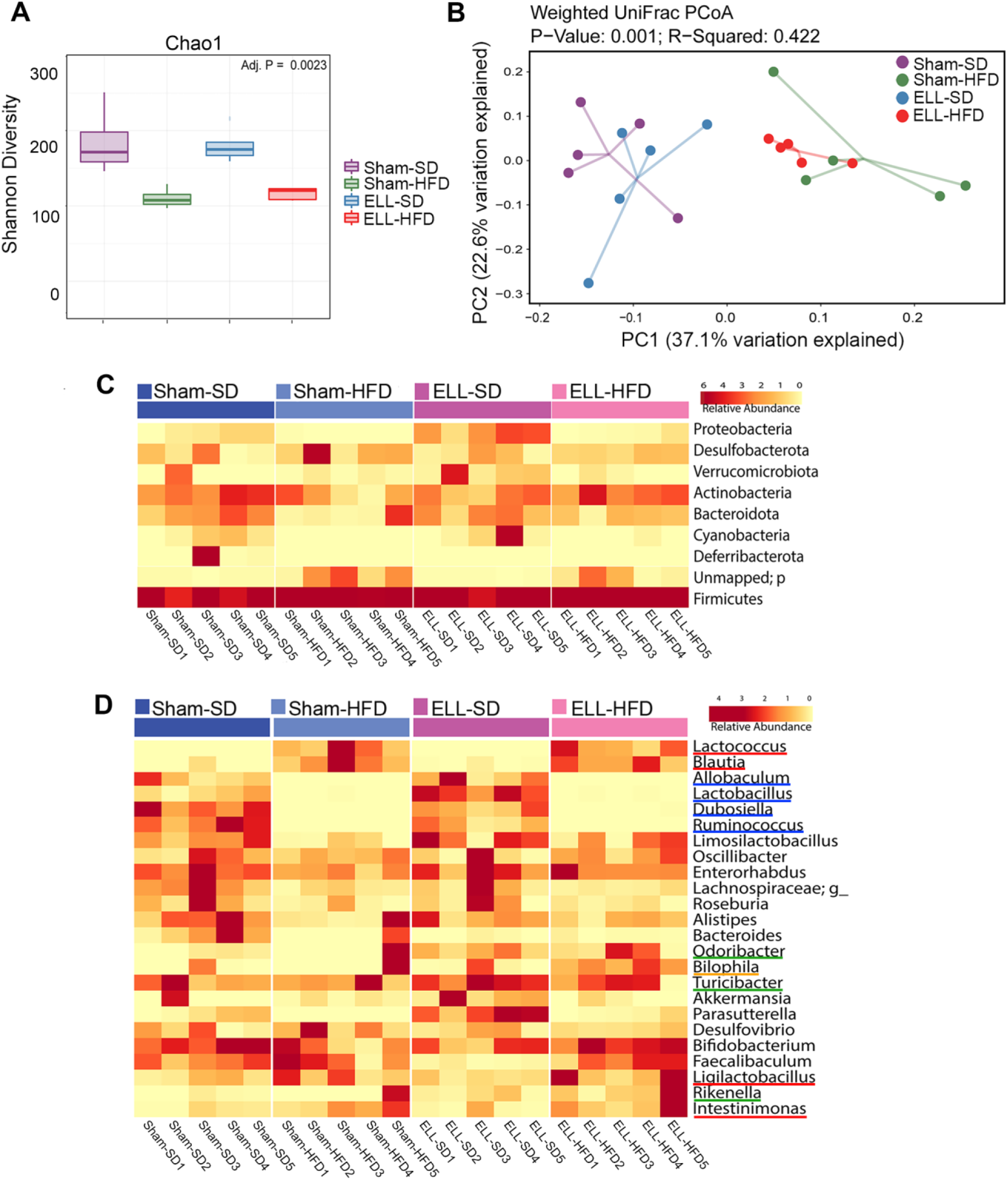
HFD altered the composition of the gut microbiota. (**A**) Box plots corresponding to the Chao1 diversity index (alpha diversity). (**B**) Principal Coordinates Analysis (PCoA) of beta-diversity based on weighted Unifrac dissimilarities in fecal samples. P = 0.001, R=0.422. (n=5 per group). (**C**) Heatmap representation of relative abundances of the phyla in feces. (**D**) Heatmap depiction of the relative abundances of the genera in feces (n=5 per group). ELL: endometrial-like lesion, SD: standard diets, and HFD: high-fat diets.

To assess whether the unique enteric bacterial profiles were attributed to specific taxa, the phyla among samples in the group were profiled (Figure 7C). The proportions of Proteobacteria and Cyanobacteria were reduced under the HFD condition, while feces in ELL-SD mice contained a higher abundance of Proteobacteria than those in Sham-SD mice. However, increased Proteobacteria were not observed in ELL-HFD mice compared to Sham-HFD mice, suggesting that the effect of HFD on Proteobacteria was stronger than lesion induction. Firmicutes and Bacteroidetes, which constitute the majority of the gut microbiota, are known to be affected by obesity, as obesity induces a reduction in the abundance of Bacteroidetes and an increase of Firmicutes proportion (49). Although the increase of Firmicutes was minor under the HFD condition, the Bacteroidetes proportion was clearly reduced in Sham-HFD mice. The Bacteroidetes population was retained in ELL-HFD mice, similar to its abundance in Sham-SD and ELL-SD mice, indicating that lesion induction increased Bacteroidetes even though mice were under the HFD condition. This result supports the study from Chadchan *et al*. that lesion induction increases the abundance of Bacteroidetes (50), while lesion induction with SD did not show a noticeable increase of Bacteroidetes in our study (Sham-SD vs ELL-SD). When we examined bacteria at the genus level, HFD clearly altered several genera among the groups (Figure 7D). In agreement with previous studies (51, 52), HFD strongly elevated *Lactococcus* and *Blautia* genera (red lines). HFD slightly increased *Ligilactobacillus* and *Intestinimonas* genera (red lines), whereas HFD mice contained negligible abundances of *Allobaculum*, *Lactobacillus*, *Dubosiella*, and *Ruminococcus* genera (blue lines). *Odoribacter*, *Turicibacter*, and *Rikenella* genera (green lines) were increased in ELL mice, and the *Bilophila* genus (orange line) was only higher in ELL-HFD mice. These data suggest that HFD or ELL alter the bacteria diversity and composition associated with endometriosis.

## Discussion

Endometriosis is generally classified into four stages according to the revised criteria from the American Society of Reproductive Medicine (rASRM) and the American Fertility Society (AFS) based on lesion size, location, and the extent of adhesions (4, 53). However, disease symptoms, such as endometriosis-associated pain, are not correlated with the staging system (4, 54). Patients with stage I disease can have severe pain, while stage IV patients can be asymptomatic (1, 55), indicating that several other factors contribute to disease symptoms. Due to the chronic inflammatory nature of endometriosis, the disease progression and symptoms can be affected by environmental factors. In the present study, our results highlight that Western-style HFD-induced obesity did not alter endometriotic lesion numbers (=disease progression) but enhanced disease-related hyperalgesia (=endometriosis-associated pain). The important findings are: 1) Peritoneal inflammatory (Ly6C+) MΦ and cytokine levels, especially TNFα, increased by lesion induction were elevated by chronic exposure to HFD. 2) Pain-related mediators, such as neurotransmitters CGRP and SP, in the DRG were further stimulated after lesion induction under the HFD condition. 3) Although HFD alone did not affect peritoneal Ly6C+ MΦ without lesion induction, the diversity and composition of the gut microbiota were clearly altered by HFD as a sign of low-grade systemic inflammation (14). Thus, HFD might not be able to establish solely a local inflammatory environment in the pelvic cavity but can contribute to further enhancing chronic inflammation associated with disease symptoms after the disease is established.

In non-human primates, rhesus macaque females exposed to testosterone (T) and/or consumed Western-style diets (WSD) at the time of menarche for 7 years developed endometriosis, especially T+WSD resulted in earlier onset of disease with high stages and large chocolate cysts (56). In a mouse model of endometriosis, HFD-induced obese mice increased lesion number and weight, which depended on leptin or leptin receptor (57). Another mouse study of endometriosis showed that HFD increased lesion number and MΦ infiltration and proinflammatory and prooxidative stress-related genes in the lesion when *Klf9* null donor endometrial fragments were inoculated as a donor tissue (58). This group further reported reduced lesion number and weight when wild-type donor tissues were used, whereas enhanced signs of inflammation were not observed in this study, indicating variability of distinct genetic dysfunctions and lesion environment for endometriosis progression (59).

One of the hallmarks of diet-induced obesity is low-grade chronic inflammation (12). Chronic consumption of HFD leads to the accumulation of MΦ and T-cells in adipose tissues to secrete proinflammatory cytokines (13). We have previously reported that lesion induction enhances the process of differentiation and maturation of monocyte-derived MΦ and increases Ly6C+ proinflammatory MΦ in the peritoneal cavity while reducing the maintenance of embryo-derived resident TIM4+ MΦ (29). The present study showed that Ly6C+ MΦ were higher in ELL mice and further increased in mice exposed to HFD, indicating the impact of HFD contribution to peritoneal inflammation after disease onset. In support of our findings, an HFD-induced proinflammatory environment promotes the differentiation of Ly6C+ monocyte into inflammatory MΦ, which migrate to the lung and worsen its pathophysiology (60). TIM4+ residential MΦ were reduced in both ELL-SD and ELL-HFD mice, whereas HFD did not further alter TIM4+ MΦ. Peritoneal inflammation can induce the macrophage disappearance reaction (MDR), by which the reduction of residential MΦ occurs. We have previously shown that extreme MDR of TIM4+ MΦ was induced 3 days after lesion induction, and it gradually recovered. However, it remains slightly diminished 6 weeks after disease onset (29). Thus, the recovery of residential TIM4+ MΦ from MDR, which includes replenishment and proliferation, is less likely affected by exposure to HFD. On the other hand, an alteration in the distribution of peritoneal T-cells by lesion induction and HFD was not observed in the study, suggesting aberrant MΦ functions might be a crucial event for establishing the chronic inflammatory state of endometriosis, as increased MΦ infiltration was also observed in the lesions under the HFD condition. However, heterogeneous T-cell functions and interaction between T-cells and MΦ remain to be studied.

In the present study, abdominal endometriosis-associated hyperalgesia was induced by lesion induction and further sensitized in ELL-HFD mice. This result was supported by the signs of sensitization of peripheral DRG, which was mediated by increased proinflammatory cytokines, TNFα, IL1β, and IL6, that are known to be increased pelvic MΦ in endometriosis patients (44) and have been targeted for pathological pain (61). Our previous studies show that PF from ELL mice stimulated DRG outgrowth, which was reduced by inhibiting cytokine and chemokine secretion in the peritoneal cavity (28). Thus, the inflammatory environment established in the pelvic cavity is critical for chronic endometriosis-associated hypersensitivity. The elevated sensitivity is not systemic, as our results showed only signs of abdominal hyperalgesia but not hind paw sensitivity by either lesion induction or HFD. Thus, it remains to study how chronic abdominal pain stimulus is delivered and maintained to the central nervous system. As endometriosis-associated pain is one of the significant problems in this disease, its mechanisms with the pathophysiology of endometriosis need to be further studied to enhance the quality of life in patients.

Our study showed that gut microbiota dysbiosis was induced by chronic exposure to HFD. HFD has been known to reduce the diversity of gut microbiota (62). The phyla Firmicutes increase while Bacteroidetes decrease, though there are variations depending on the differences in diet compositions and exposure duration (63, 64). Interestingly, our results showed a lower abundance of Proteobacteria in HFD mice, whereas increased Proteobacteria abundance with HFD consumption has been reported (65). Increased *Allobaculum* abundance has been shown under the HFD condition (66), though the abundance of *Allobaculum* was reduced in our HFD mice. However, this inconsistency is likely due to different types of diet, fat, and other environmental factors in the various studies (14). Despite having variable alterations of gut microbiota, HFD-induced dysbiosis increases gut permeability and creates chronic inflammation, affecting inflammatory diseases directly or indirectly (67).

The present study shows endometriosis-associated abdominal hyperalgesia was escalated under exposure to HFD, whereas lesion progression was not altered. These results include increased proinflammatory MΦ and cytokine levels in the peritoneal cavity, neuromodulators in the DRG, and dysbiosis of gut microbiota. Retrograde menstruation causes massive inflammatory responses in the pelvic cavity, which involves the recruitment of monocytes that differentiate into proinflammatory MΦ and secrete cytokines and chemokines (27). However, the acute inflammation associated with retrograde menstruation typically resolves by the next menstrual cycle. If women are under systemic low-grade inflammation induced by environmental factors like HFD, it is expected to be hard to solve this acute incidence. As menstrual cycles repeatedly occur in women, each retrograde menstruation induces composite inflammation in the pelvic cavity, and unsolved inflammation can worsen to develop chronic conditions further. Thus, the present results suggest that diet-induced obesity could be a risk factor for establishing a chronic inflammatory environment and severe endometriosis-associated pain, which can be independent of disease progression.

## Materials and Methods

### Animals

All animal experiments were performed at Washington State University according to the NIH guidelines for the care and use of laboratory animals (protocol #6751). C57BL/6 (Jax #664) mice were obtained from the Jackson Laboratory and used for the studies.

### Mouse Model of Endometriosis

An experimental mouse model of endometriosis was established by adopting procedures described previously (28, 29, 48, 68–70). Briefly, a ‘menses-like’ event was induced in ovariectomized estradiol-17β (E_2_)- and progesterone-primed donor mice following an established protocol (71). Then, mouse menses-like endometrium scraped from myometrium and cut into fragments (1-2 mm per side) were introduced as the source of syngeneic mouse endometrium (donor) via injection (in 0.2 mL PBS) into the peritoneal cavity of ovariectomized E_2_-primed mice (recipient) under anesthesia via inhaled isoflurane.

### Study Design

To induce diet-dependent obese mice, female mice were fed Teklad Rodent Diet (#2019, Envigo) as SD (Washington State University regular diet) that contain 9% of total calories from fat or HFD (D12451, Research Diets) that contain 45% of total calories from fat starting at the age of 5 weeks (defined as Week 0 of the 18-week program, Figure 1A). BW was recorded once a week. Twelve weeks after SD or HFD feeding, mice were further assigned to sham control without lesion induction or ELL-induced groups. Thus, there were a total of 4 groups with Sham-SD (n=8), Sham-HFD (n=8), ELL-SD (n=18) and ELL-HFD (n=18). Six weeks after induction, a behavior test was performed, and fresh feces were collected and immediately frozen at -80C. Mice were then euthanized for sample collections: blood was collected via cardiac puncture, PF was recovered by lavage (4 mL x 2 of ice-cold PBS), and ELL and bilateral lumbar (L4-6) DRG were collected for further analysis. To obtain baseline results of behavior and peritoneal lavage, a separate pool of non-induced mice was fed SD or HFD for 12 weeks and used for behavior testing (n=10) and sample collection (n=5). Blood glucose levels (n=5) were measured by Contour Next (Ascensia Diabetes Care), and plasma insulin (n=5) was analyzed by ELISA (EZRMI-13K, Sigma Aldrich), according to the manufacturer’s instructions.

### Von Frey test

A behavioral (mechanical sensitivity) test was performed before sample collection. Mice were allowed to acclimate in the testing room for 30 min, and then the von Frey test was performed using von Frey Filaments (BIO-VF-M, Bioseb). Filaments were applied 10 times to the skin perpendicular to the lower abdomen and bilateral hind paws. The force in grams (g) of the filament evoking a withdrawal response (50% response count as sensitive) was recorded. Three behaviors were considered positive responses to filament stimulation: 1) sharp retraction of the abdomen, 2) immediate licking and/or scratching of the area of filament stimulation, or 3) jumping. All behavioral tests were blind to investigators. Mice without ELL or sham induction after 12 weeks of SD or HFD feeding were included as a baseline result.

### Flow cytometry

Peritoneal lavages were centrifuged to collect peritoneal exudate cells. After lysing red blood cells by 1x RBC Lysis Buffer (BioLegend), cells were incubated at room temperature for 20 minutes with Zombie Aqua™ Fixable Viability dye (BioLegend) and blocked on ice for 20 minutes with Fc Block anti-CD16/CD32 (Thermo Fisher). Then, cells were stained with fluorochrome-conjugated monoclonal antibodies (Supplementary Table S1) for 1 hour. Samples were acquired with the Attune NxT Acoustic Focusing Cytometer using Attune NxT software (Thermo Fisher), and data were analyzed with FlowJo v10.9. For analysis, only singlets (determined by forward scatter height vs. area) and live cells (Zombie Aqua negative) were used.

### Immunofluorescence

Immunostaining of BDNF, CGRP, SP, TRPV1, neurofliment (NF), and CD68 was performed with cross-sections (5 μm) of paraffin-embedded tissues using specific commercially available primary antibodies (Supplementary Table S1) and AlexaFluor 488 and 568-conjugated F(ab’) secondary antibody (Molecular Probe) or VECTASTAIN ABC kit (Vector lab). Cell-specific CD68 positive and total cell numbers were counted in the area of 0.07244 mm^2^, and the percentage of CD68+ cells was shown. NF was used as a pan-neuronal marker and was co-stained with BDNF, CGRP, SP, or TRPV1. BDNF, CGRP, SP, or TRPV1 positive cells in the DRG were counted in the area of 0.07244 mm^2^. The percentages of BDNF, SP, CGRP, or TRPV1 positive cells per NF-positive DRG were shown.

### IQELISA

Protein yield from PF was quantitated by BCA assay (Pierce), and TNFα (IQM-TNFA-1), IL1β (IQM-IL1b-1), IL6 (IQM-IL6-1), and IL10 (IQM-IL10-1) were further quantified by IQELISA kits (Ray Biotech) according to the manufacturer’s instructions.

### Bacterial 16S rRNA gene sequencing and analysis

DNA was extracted from fecal pellets (100 mg, n=5 per group) by the QIAmp Power Fecal DNA kit (12850-50, Qiagen). The V4 region of 16S rRNA gene was amplified, and sequencing was performed on an Illumina platform by the Alkek Center for Metagenomics and Microbiome Research at Baylor College of Medicine. Demultiplexed reads were quality filtered after initial trimming, and taxonomic information was retrieved by mapping against SILVA version 138.1 (72) using an identity threshold of 70% in Quantitative Insights Into Microbial Ecology (73). Raw data in FASTQ format were uploaded to the NCBI Sequence Read Archive (PRJNA1007658). This dataset was used for downstream alpha and beta diversity analysis, and top taxa were identified using a mean abundance threshold of ≥ 0.05, as described previously (50). The alpha diversity was measured using Chao1 distances, while the beta diversity was estimated using weighted UniFrac measures (74).

### Statistical analysis

Data at 18 weeks were subjected to one-way ANOVA and Tukey’s post hoc test to identify differences among the groups using Prism software (Ver. 9.1.0, GraphPad). Data at 12 weeks and lesions at 18 weeks were analyzed by two-tailed Student’s t-test comparing SD and HFD. All experimental data are presented as mean with standard error of the mean (SEM). The Mann-Whitney test was used to compare alpha diversities among the groups. Principal Coordinates Analysis (PCoA) based on unweighted UniFrac distances were used to measure the beta diversity. Unless otherwise indicated, a *P* value less than 0.05 was considered to be statistically significant.

## Study approval

The use of animal protocol (#6751) in this study was reviewed and approved by the Institutional Animal Care and Use Committees (IACUCs) of Washington State University.

## Data availability

Raw data for bacterial 16S rRNA gene sequencing are available in the NCBI Sequence Read Archive under accession number PRJNA1007658.

## Author contributions

M.S. and K.H. designed research; T.H.W., M.S., and M.E.H. conducted animal study and sample collection; T.H.W. and M.E.H. performed ELL induction; T.H.W. performed immunofluorescence and IQELISA; M.S. performed immunofluorescence and flow cytometry; M.S. and M.E.H. performed the von Frey test; T.H.W., M.S., C.T., R.K., J.A.M., and K.H. analyzed data; K.H. wrote the paper; all authors read, reviewed and approved the manuscript.

## Acknowledgments

This work was supported by NIH/NICHD, R01HD104619 (to K.H.) and R01HD102680 (to R.K.). We thank Logan C. Butler for feeding diets and caring for mice.

## Conflict of Interest

The authors declare that they have no competing interests.

---

**Supplementary Figure S1.**
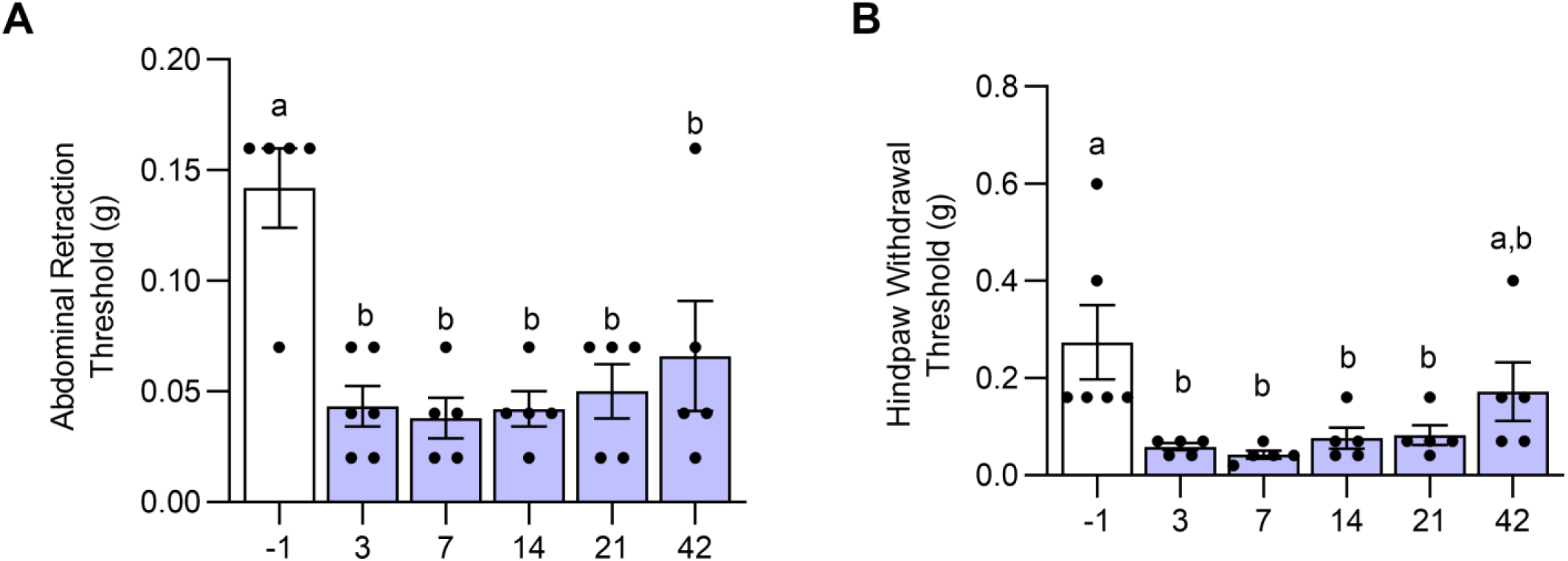
Evaluation of time-dependent endometriosis-associated hyperalgesia after ELL induction. (**A**) Abdominal retraction and (**B**) hind paw withdrawal thresholds were evaluated by the von Frey test with SD on Days -1, 3, 7, 14, 21, and 42 after ELL induction. Data are shown as mean ± SEM (n = 5). One-way ANOVA followed by Tukey’s post hoc test was used to compare the differences in abdominal and hind paw withdrawal thresholds across time points. a vs b; P<0.05. ELL: endometrial-like lesion.

**Supplementary Figure S2.**
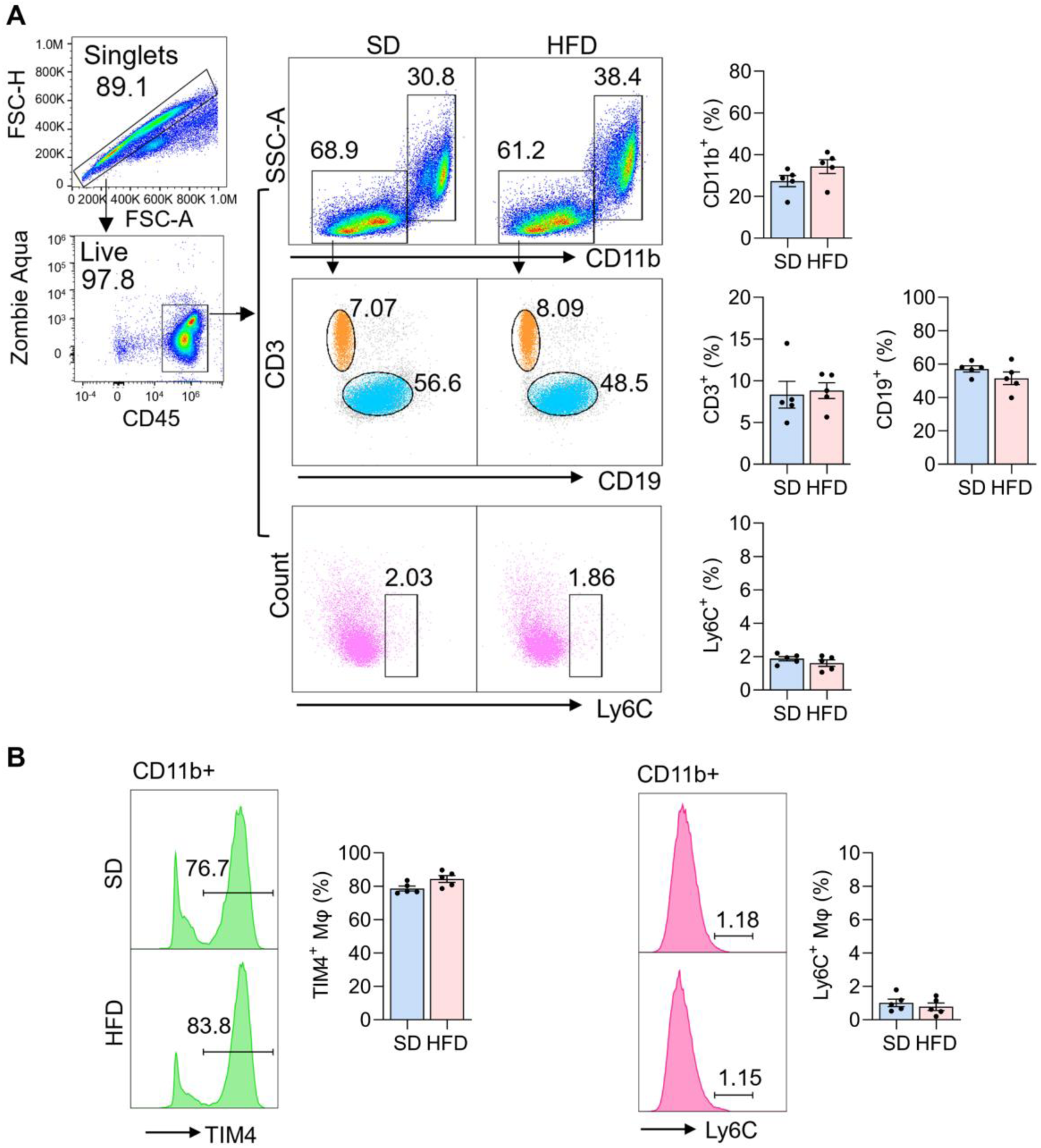
Flow cytometer analysis for peritoneal immune cells at 12 weeks (pre-induction stage). (**A**) Quantification of CD11b+ (MΦ), CD3+ (T-cells), CD19+ (B-cells), and Ly6C+ cells (n=5) (**B**) TIM4+ and Ly6C+ MΦ were quantified in the PF (n=5). Student t-test was used to compare the difference between SD and HFD groups (no difference was detected). SD: standard diets, HFD: high-fat diets, and MΦ, macrophages.

**Table S1.**
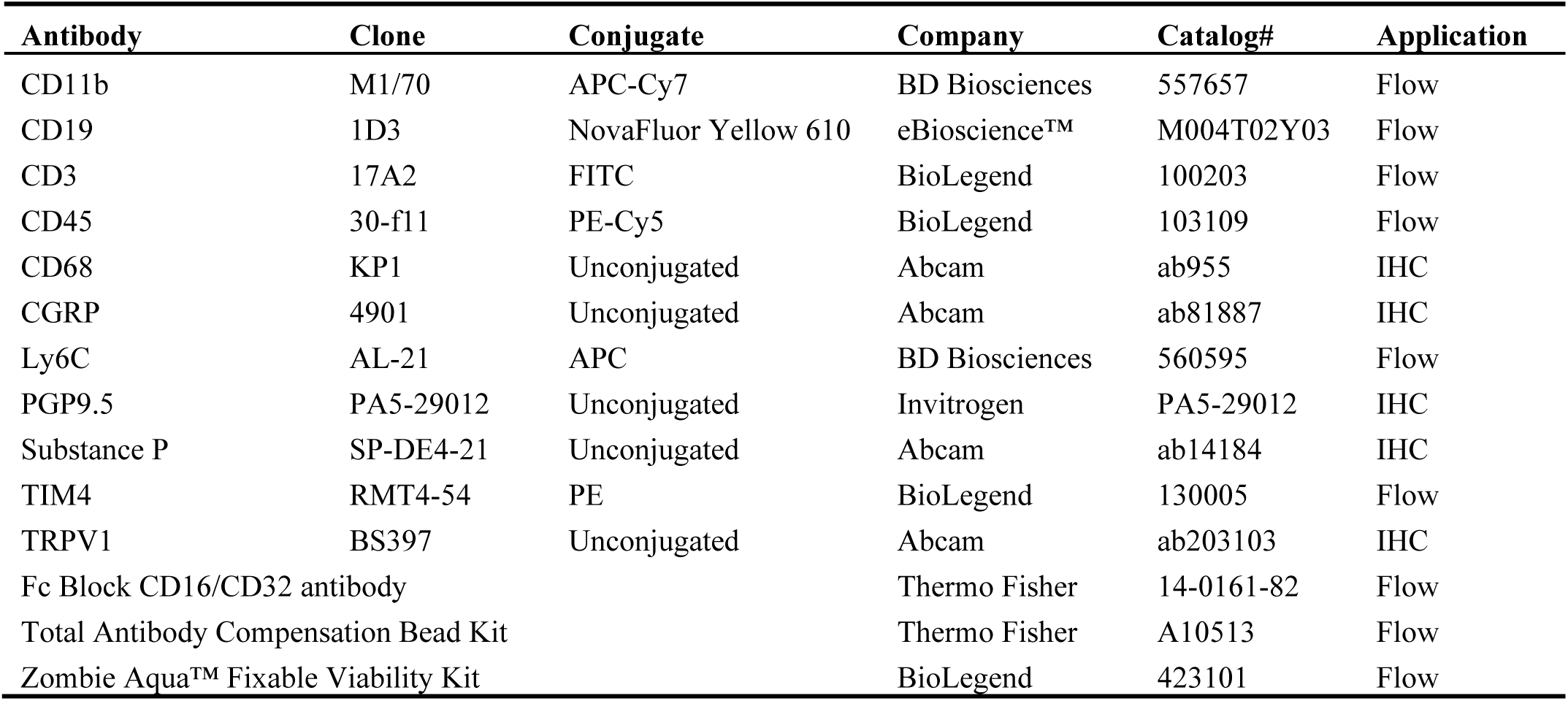
Antibodies and Reagents for Flow Cytometry and Immunochemistry.

## Notes

### Competing Interest Statement

The authors have declared no competing interest.

