## Supplementary material for "High-fat diets promote peritoneal inflammation and augment endometriosis-associated abdominal hyperalgesia": Suppl Table and Figures

### Table S1 Antibodies and Reagents for Flow Cytometry and Immunochemistry

| Antibody | Clone | Conjugate | Company | Catalog# | Application |
|---|---|---|---|---|---|
| CD11b | M1/70 | APC-Cy7 | BD Biosciences | 557657 | Flow |
| CD19 | 1D3 | NovaFluor Yellow 610 | eBioscience™ | M004T02Y03 | Flow |
| CD3 | 17A2 | FITC | BioLegend | 100203 | Flow |
| CD45 | 30-f11 | PE-Cy5 | BioLegend | 103109 | Flow |
| CD68 | KP1 | Unconjugated | Abcam | ab955 | IHC |
| CGRP | 4901 | Unconjugated | Abcam | ab81887 | IHC |
| Ly6C | AL-21 | APC | BD Biosciences | 560595 | Flow |
| PGP9.5 | PA5-29012 | Unconjugated | Invitrogen | PA5-29012 | IHC |
| Substance P | SP-DE4-21 | Unconjugated | Abcam | ab14184 | IHC |
| TIM4 | RMT4-54 | PE | BioLegend | 130005 | Flow |
| TRPV1 | BS397 | Unconjugated | Abcam | ab203103 | IHC |
| Fc Block CD16/CD32 antibody | | | Thermo Fisher | 14-0161-82 | Flow |
| Total Antibody Compensation Bead Kit | | | Thermo Fisher | A10513 | Flow |
| Zombie Aqua™ Fixable Viability Kit | | | BioLegend | 423101 | Flow |

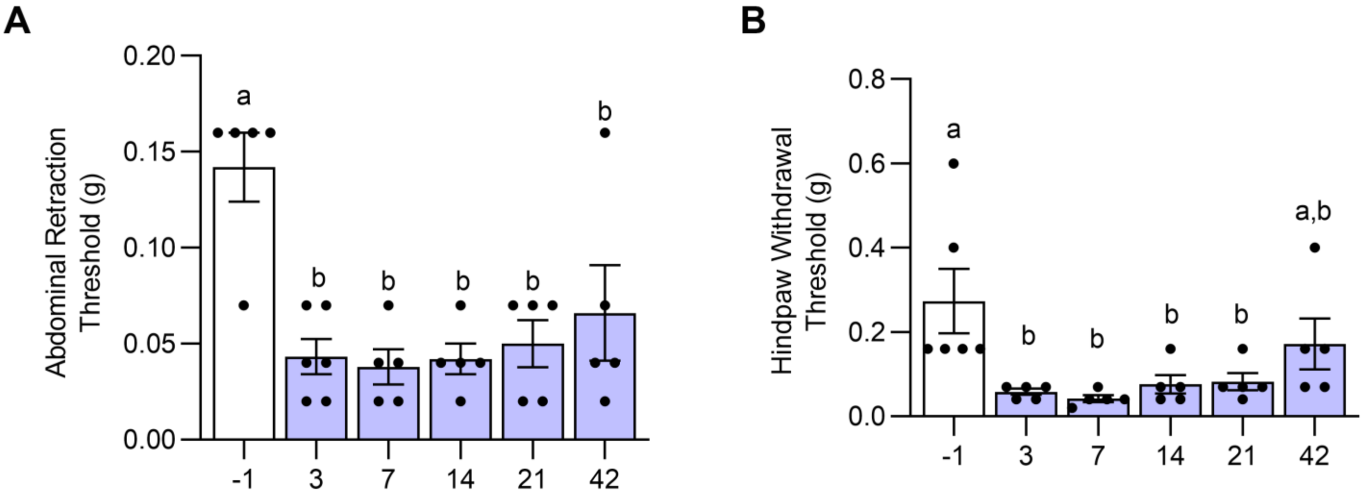


**Supplementary Figure S1**. Evaluation of time-dependent endometriosis-associated hyperalgesia after ELL induction. (A) Abdominal retraction and (B) hind paw withdrawal thresholds were evaluated by the von Frey test with SD on Days -1, 3, 7, 14, 21, and 42 after ELL induction. Data are shown as mean ± SEM (n = 5). One-way ANOVA followed by Tukey's post hoc test was used to compare the differences in abdominal and hind paw withdrawal thresholds across time points. a vs b; $P<0.05$. ELL: endometrial-like lesion.

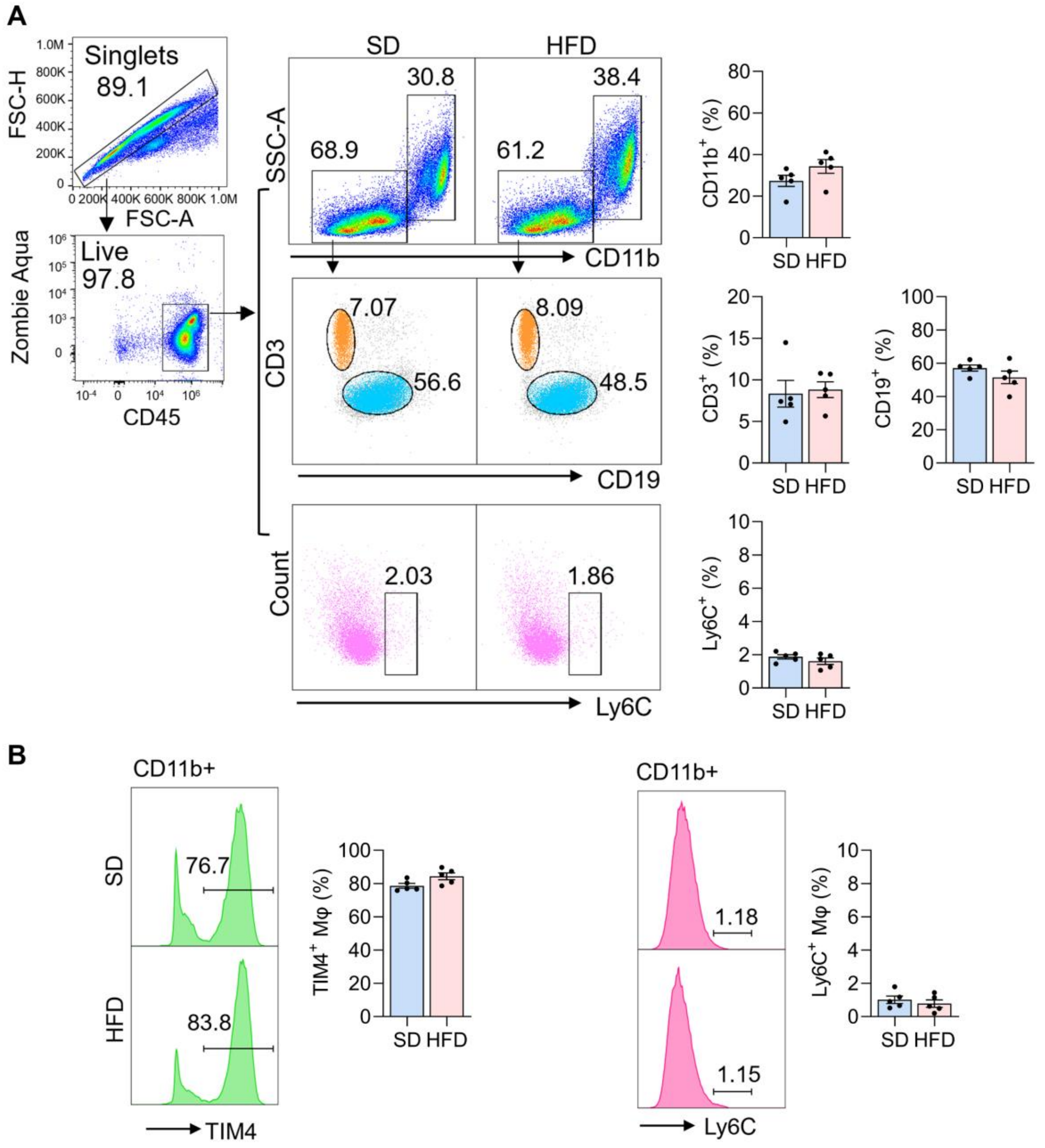


**Supplementary Figure S2.** Flow cytometer analysis for peritoneal immune cells at 12 weeks (pre-induction stage). (A) Quantification of CD11b+ (MΦ), CD3+ (T-cells), CD19+ (B-cells), and Ly6C+ cells (n=5) (B) TIM4+ and Ly6C+ MΦ were quantified in the PF (n=5). Student t-test was used to compare the difference between SD and HFD groups (no difference was detected). SD: standard diets, HFD: high-fat diets, and MΦ, macrophages.
